# CD34-positive monocytes are highly susceptible to HIV-1

**DOI:** 10.1101/2024.02.26.582226

**Authors:** Naofumi Takahashi, Osamu Noyori, Yoshihiro Komohara, Youssef M. Eltalkhawy, Masatoshi Hirayama, Ryoji Yoshida, Hideki Nakayama, Marcelo J. Kuroda, Takushi Nomura, Hiroshi Ishii, Tetsuro Matano, Hiroyuki Gatanaga, Shinichi Oka, Masafumi Takiguchi, Shinya Suzu

## Abstract

HIV-1 persists in cellular reservoirs despite effective combined antiretroviral therapy (cART). CD4^+^ T cells are a well-known reservoir, but there is evidence suggesting that myeloid cells, including circulating monocytes, are also a clinically relevant reservoir. However, it is not fully understood which subsets of monocytes are preferentially infected in vivo. Here, we show that a monocyte fraction expressing a stem cell marker CD34 is more susceptible to HIV-1 infection than the CD34-negative major subset. In cART-untreated viremic individuals, the CD34^+^ fraction increased in the percentage in total monocytes, and harbored higher copies of proviral DNA than the major subset. Consistent with this, the CD34^+^ fraction expressed HIV-1 receptors CD4 and CCR5 at higher levels and HIV-1 restriction factors MX2 and SAMHD1 at lower levels. Interestingly, proviral DNA was still detectable in the CD34^+^ fraction of cART-treated virologically suppressed individuals. CD34^+^ monocytes were also present in lymph nodes, and expressed CD4 and CCR5 at higher levels than the major subset, as observed in peripheral blood. Moreover, CD34^+^ monocytes present in peripheral blood and lymph nodes highly expressed CCR7 and sphingosine-1-phosphate receptor 1 (S1PR1), critical regulators of in vivo cellular trafficking. Collectively, our findings raise the new possibility that lymph node CD34^+^ monocytes, which originate from the circulation, are infected with HIV-1 owing to their high susceptibility to HIV-1, and return to circulation, which explains the detection of proviral DNA in peripheral CD34^+^ monocytes even after long-term cART.

## Introduction

Antiretroviral therapy (cART) suppresses HIV-1 to undetectable levels in plasma. However, an obstacle to cure is that HIV-1 integrates into the genome of infected cells and persists in cellular reservoirs, and therefore grows once cART is interrupted. Resting CD4^+^ T cells are the well-known reservoir (1). Meanwhile, there is growing evidence that myeloid cells, such as tissue macrophages and circulating monocytes, can also serve as reservoirs (2, 3). For instance, HIV-1 particles carrying myeloid-specific markers on their surface were detected in plasma of individuals undergoing analytical cART interruption (4), suggesting that these particles were produced by myeloid cells. Indeed, HIV-1-infected macrophages have been detected in several tissues of individuals on cART (5, 6).

A recent study also emphasized the importance of monocytes as the long-lived latently or persistently infected cells towards HIV-1 cure. Veenhuis *et al.* detected intact proviral DNAs in monocytes of individuals on long-term cART, and demonstrated that viruses produced by ex vivo-activated monocytes could infect bystander T cells, resulting in viral spread (7). Although most proviruses in monocytes were defective as observed in CD4^+^ T cells (7), these cells harboring defective proviruses are another important target towards HIV-1 cure since RNAs and proteins are often expressed from the defective proviruses, which could alter anti-viral T cell responses and myeloid cell functions that contribute to HIV-1-associated chronic comorbidities (8). However, it is not fully understood which subsets of monocytes are preferentially infected in vivo.

Monocytes consist of at least three subsets: classical CD14^+^CD16^−^, intermediate CD14^+^CD16^+^, and nonclassical CD14^−^CD16^+^ (9). Classical monocytes are the major subset constituting 80-85% of the total population, with a small fraction expressing a hematopoietic stem cell marker CD34 (10–14). The CD34^+^ fraction (CD14^+^CD16^−^CD34^+^) has been reported to contain fibroblastic leukocytes called fibrocytes (12), and exhibits features of stem cells (13) or regulatory monocytes (14). A similar CD34^+^ fraction has been detected in adipose tissues (15, 16). However, the precise role of this fraction remains largely unknown. Our previous analysis of cART-untreated viremic individuals revealed that circulating CD14^+^CD16^−^CD34^+^ monocytes contained more HIV-1 proviral DNA than CD34-negative monocytes (17), suggesting that the CD34^+^ fraction was highly infected with HIV-1 in the major subset. In this study, we therefore sought to clarify how and why the CD34^+^ monocytes are susceptible to HIV-1 infection, and whether the CD34^+^ monocytes harbor proviral DNA in virally suppressed individuals.

## Materials and Methods

### Sorting and proviral PCR of peripheral blood cells of HIV-1-infected individuals

cART-naive Japanese individuals chronically infected with HIV-1 and cART-treated (>3 years) individuals were recruited from the National Center for Global Health and Medicine, Japan. This study was approved by the Ethics Committees of the National Center for Global Health and Medicine and Kumamoto University, Japan. Heparinized venous blood was collected from these individuals after informed consent was obtained in accordance with the Declaration of Helsinki. To sort CD34^+^ monocytes or CD34^−^ monocytes, peripheral blood mononuclear cells (PBMCs) were stained with the following antibodies: allophycocyanin (APC)-anti-CD34 (581; BioLegend), FITC-anti-CD14 (61D3; eBioscience), Pacific Blue-anti-CD16 (3G8; BioLegend), and PE-anti-CD3 (OKT3; BioLegend). To sort resting CD4^+^ T cells, PBMCs were stained with APC-anti-CD4 (RPA-T4; BioLegend), FITC-anti-CD25 (M-A251; BioLegend), Pacific Blue-anti-CD69 (FN50; BioLegend), and PE-anti-HLA-DR (LN3; BioLegend). The cells in the live cell gates were sorted using FACSAria or FACSAria Ⅱ (BD Biosciences), as described previously (17). CD34^+^ monocytes (CD3^−^CD14^+^CD16^−^CD34^+^), CD34^−^ monocytes (CD3^−^CD14^+^CD16^−^CD34^−^), or resting CD4^+^ T cells (CD4^+^CD25^−^CD69^−^HLA-DR^−^) were sorted into 96-well plates (2,000 cells/well), and genomic DNA was isolated using QIAamp DNA micro kit (QIAGEN). Two-step PCR to amplify *gag* or *pol* region was performed with eight different primer pairs (Supplemental Table 1) and Ex-Taq (TaKaRa-Bio) as described previously (17). The first PCR was performed using genome DNA (100 cells/reaction, 38 cycles, the initial denaturation at 98°C for 2 min, 98°C for 10 sec, 55°C for 30 sec and 72°C for 1 min, and a final extension at 72°C for 7 min), and the second PCR was performed using an aliquot of the first PCR products (1/2,500 dilution, 38 cycles). As a loading control, GAPDH was amplified (35 cycles) with a primer pair 5’-CCACCCTGTTGCTGTAGCCAAATTCG-3’ and 5’-TCCGGGAAACTGTGGCGTGATGG-3’.

### Human lymph node cells

We also analyzed regional lymph nodes resected from patients with oral squamous cell carcinoma who underwent radical resection at Kumamoto University. This study was conducted with the approval of the Ethics Committees of Kumamoto University, and in accordance with the guidelines for Good Clinical Practice and the Declaration of Helsinki. Lymph nodes collected mainly from levels Ⅰ through Ⅲ were minced with tweezers and scalpels, and incubated with collagenase D (1 mg/ml; Roche) in RPMI1640 medium at 37 °C for 30 min. Cells were filtered using a 75-μm cell strainer, and subjected to flow cytometric analysis.

### Flow cytometry

CD34^+^ monocytes in peripheral blood or lymph nodes were characterized by flow cytometry. Cells were stained with fluorescent dye-labeled antibodies after incubation with Fc block reagent (BD Biosciences), and analyzed on the FACSCanto Ⅱ (BD Biosciences) using the FlowJo software (Tree Star). The antibodies used were as follows: APC-anti-CD34, FITC-anti-CD14, Pacific Blue-anti-CD14 (M5E2; BioLegend), peridinin chlorophyll protein-cyanin 5.5 (PerCP-Cy5.5)-anti-CD16 (3G8; BioLegend), Pacific Blue-anti-CD45 (HI30; BioLegend), PerCP-Cy5.5-anti-CD45 (HI30; BioLegend), PE-anti-CD3 (OKT3; BioLegend), PE-anti-CD19 (HIB19; BioLegend), APC-anti-CD64 (10.1; BioLegend), PerCP-Cy5.5-anti-CD163 (GHI/61; BioLegend), PE-Cyanine7 (PE-Cy7)-anti-CCR2 (K036C2; BioLegend), APC-anti-HLA-A/B/C (W6/32; BioLegend), PE-anti-CD4 (RPA-T4; BioLegend), APC-Cy7-anti-CD4 (A161A1; BioLegend), FITC-anti-CCR5 (HEK/1/85a; BioLegend), PE-anti-SLAMF7 (162.1; BioLegend), PE-Cy7-anti-CCR7 (G043H7; BioLegend), and Alexa Fluor 488-anti-S1PR1 (218713; R&D Systems).

### Quantitative RT-PCR

CD34^+^ monocytes (CD3^−^CD14^+^CD16^−^CD34^+^) or CD34^−^ monocytes (CD3^−^CD14^+^CD16^−^CD34^−^) of healthy donors were purified by sorting, and cellular RNA was isolated using RNeasy micro kit (QIAGEN). Complementary DNA was prepared using M-MLV RT (Invitrogen), and quantitative PCR (qPCR) was performed with SYBR Premix Ex Taq II (TaKaRa-Bio) using a LightCycler (Roche) as described previously (18). The expression levels of mRNA of HIV-1 restriction factors (MX2, SAMHD1, BST2, and APOBEC3G) were calculated using the ΔΔCt method and normalized to that of GAPDH. The sequences of the primer pairs used were as follows: 5’-TTACTGAAATAGGCATCCACCTGA-3’ and 5’-TAAAATACTGAATTATAAATGGGATCTGGT-3’ (MX2), 5’-GGATTACTAAAAACCAGGTTTCACAACT-3’ and GCTCTGCAAATTTCTCTGGCAG-3’ (SAMHD1), 5’-TTCTCAGTCGCTCCACCT-3’ and 5’-CACCTGCAACCACACTGT-3’ (BST2), 5’-GGCTCCACATAAACACGGTTTC-3’ and 5’-AAGGGAATCACGTCCAGGAA-3’ (APOBEC3G), and 5’-GACTCATGACCACAGTCCATGC-3’ and 5’-GAGGAGACCACCTGGTGCTCAG-3’ (GAPDH).

### Microarray

CD34^+^ monocytes or CD34^−^ monocytes of a healthy donor were purified by sorting, and cellular RNA was isolated using RNeasy micro kit, which were biotin-labeled using a GeneChip 3’ IVT express kit (Affymetrix). Microarray analysis was performed at TaKaRa-Bio using high-density oligonucleotide array (Human Genome U133 Plus 2.0) and GeneSpring 12.5 software (Agilent Technologies) as described previously (17). The data have been deposited in the National Center for Biotechnology Information Gene Expression Omnibus (GSE229346; http://www.ncbi.nlm.nih.gov/geo).

### Macrophage differentiation

CD34^+^ monocytes or CD34^−^ monocytes of healthy donors were purified by sorting, and cultured with RPMI1640-10% FCS containing 100 ng/ml recombinant human (rh)M-CSF (a gift from Morinaga Milk Industry) for 7–10 days to induce their differentiation into macrophages (17). The adherent cells were detached using enzyme-free cell-dissociation buffer (Gibco) and subjected to flow cytometric analysis. The antibodies used were as follows: Pacific Blue-anti-CD14 and PE-anti-CD163 (GHI/61; BioLegend).

### Lymph node cells of rhesus macaques

We also analyzed rhesus macaques infected with HIV-1-related simian immunodeficiency virus (SIV). This study was approved by the Tulane University Institutional Animal Care and Use Committee and performed in accordance with the NIH Guide for the Care and Use of Laboratory Animals (19), or approved by the Committee on the Ethics of Animal Experiments of the Institute for Virus Research, Kyoto University and National Institutes of Biomedical Innovation, Health and Nutrition and performed in accordance with the Guidelines for Proper Conduct of Animal Experiments established by Science Council of Japan (20). Lymph nodes collected from cART-untreated chronically SIV-infected rhesus macaques were subjected to several analyses. Immunohistochemical analysis was performed essentially as described previously (21). Antibodies used were anti-CD14 (4B4F12; Abcam) and anti-CD8 (C8; Nichirei, Tokyo, Japan). Secondary antibodies were purchased from Nichirei, and reactions were visualized using the diaminobenzidine substrate system (Nichirei). Cells in lymph nodes (CD45^+^CD3^−^CD14^+^CD34^+^ or CD45^+^CD3^+^CD19^−^CD3^−^) were sorted using FACSAria III and analyzed for proviral DNA. Antibodies used were as follows: PerCP-anti-CD45 (D058-1283; BD Biosciences), APC-Cy7-anti-CD3 (SP34-2; BD Biosciences), PE-Cy7-anti-CD14 (M5E2; BioLegend), PE-anti-CD34 (581; BioLegend), Pacific Blue-anti-CD19 (HIB19; BioLegend), Pacific Blue-anti-CD8 (RPA-T8; BioLegend). Two step PCR to amplify *gag* region was performed using KOD FX Neo (TOYOBO) and primer pairs 5’-AGAAACTCCGTCTTGTCAGG-3’ and 5’-TGATAATCTGCATAGCCGC-3’ (first PCR, 40 cycles, the initial denaturation at 94°C for 2 min, 98°C for 10 sec, 60°C for 30 sec and 68°C for 37 sec), 5’-GATTAGCAGAAAGCCTGTTGG-3’ and TGCAACCTTCTGACAGTGCG-3’ (second PCR, the initial denaturation at 94°C for 2 min, 98°C for 10 sec, 62°C for 30 sec and 68°C for 25 sec). In addition, the intracellular expression of viral p27 Gag proteins was assessed by flow cytometry with a CytofixCytoperm kit (BD Biosciences). Antibodies used were as follows: PerCP-anti-CD45, APC-Cy7-anti-CD3, PE-Cy7-anti-CD14, PE-anti-CD34, and Alexa Fluor 647-anti-p27 Gag (Advanced Bioscience Laboratories).

### Statistical analysis

The statistical significance of inter-sample differences was determined using a paired Student’s *t*-test. Statistical significance was set at *p* < 0.05.

## Results

### CD34^+^ monocytes are more susceptible to HIV-1 infection than CD34-negative monocytes in vivo, and show proviral persistence

CD34^+^ cells have been identified in CD14^+^CD16^−^ classical monocytes as a minor fraction (10–14) (Fig. 1A). In this study, we initially found that when compared to uninfected control, cART-untreated HIV-1-infected individuals showed an increased percentage of the CD34^+^ fraction in total classical monocyte population (Fig. 1B). As reported (17), the CD34^+^ fraction more frequently harbored HIV-1 proviral DNA than CD34-negative fraction (Fig. 1C). In the two-step proviral PCR, eight different primer pairs were used to cover a broad area of *gag* or *pol* region (Supplemental Table 1), and the positive PCR were plotted in red (Fig. 1C). This semi-quantitative PCR (100 cells/reaction in the first PCR) was reproducible since samples collected from the same individual but different visits with different plasma viral load (PVL; HIV-1 RNA copies/ml) showed the same pattern (KI110-1 and KI110-2; Fig. 1C). In this study, we also analyzed individuals treated with cART for more than 3 years (3 – 19 years, median=12.7 years) and detected proviral DNA in the CD34^+^ fraction of several individuals (Fig. 1D). Plasma viral load of these individuals was below the detection limit (<20 copies/ml) with the exception of an individual (KU09 in group-ⅲ). On a per-cell basis, the frequency of proviral DNA in CD34^+^ monocytes (CD3^−^CD14^+^CD16^−^CD34^+^) was similar to that of resting CD4^+^ T cells (CD4^+^CD25^−^CD69^−^HLA-DR^−^) (Fig. 1D). In detail, both were positive in group-ⅰ (n=9), both were negative in group-ⅱ (n=3), only resting CD4^+^ T cells were positive in group-ⅲ (n=2), and only CD34^+^ monocytes were positive in group-ⅳ (n=2). Although PCR reaction containing genomic DNA from a four-fold higher number of resting CD4^+^ T cells (i.e., 400 cells/reaction in the first PCR) was also tested for the two individuals in group-ⅳ (KU04 and KU06), we did not detect proviral DNA under the conditions (data not shown), supporting the idea that the proviruses detected in the CD34^+^ monocyte fraction were not due to contaminated resting CD4^+^ T cells, if any. These results suggest that CD34^+^ monocytes are more susceptible to HIV-1 infection than CD34-negative monocytes in vivo, which may relate to the long-term proviral persistence in CD34^+^ monocytes.

**Figure 1.**
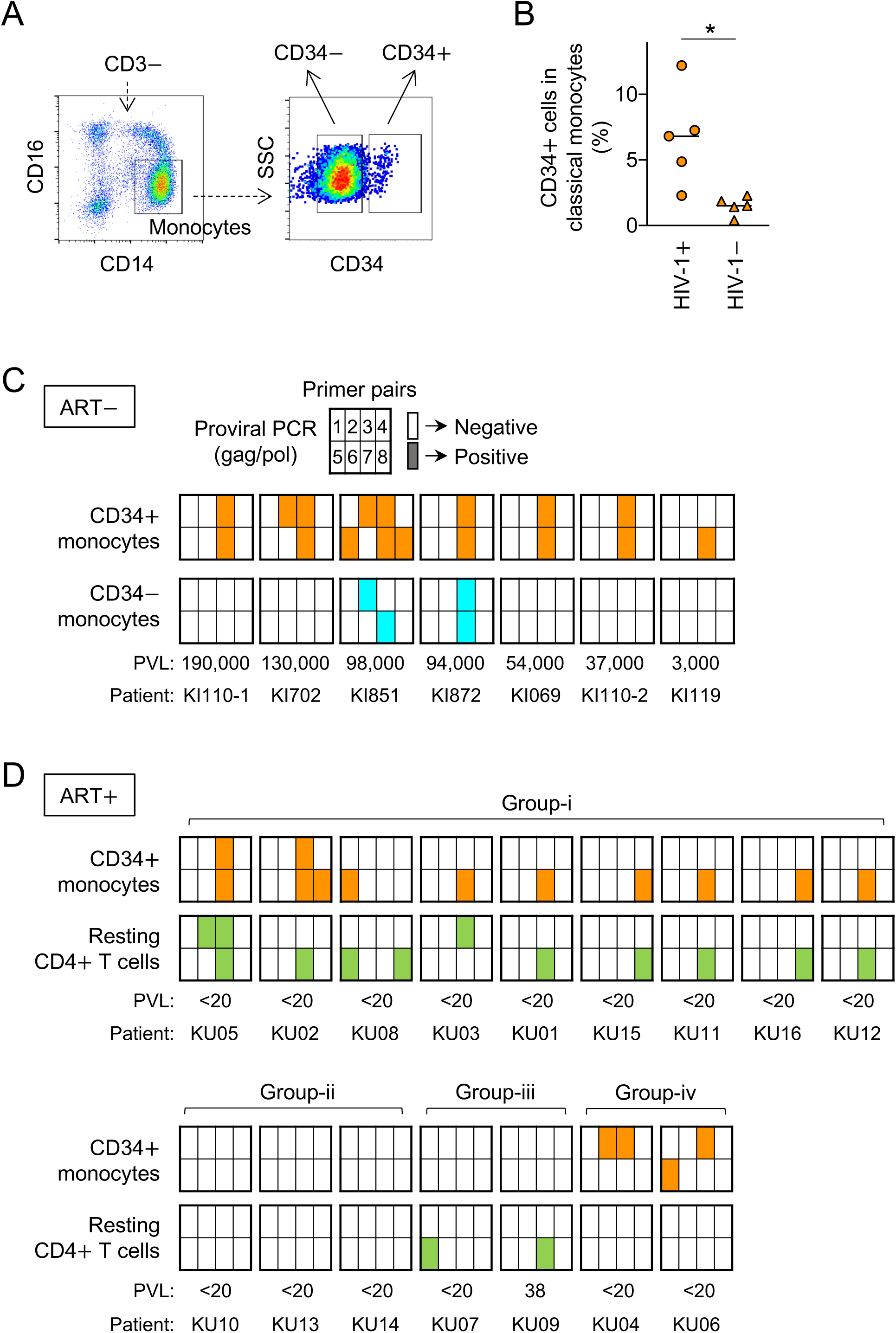
HIV-1 proviral DNA in peripheral CD34^+^ monocytes. (A) Peripheral CD34^+^ monocytes (“CD34+”, CD3^−^CD14^+^CD16^−^CD34^+^) and CD34^−^ monocytes (“CD34−”, CD3^+^CD14^+^CD16^−^CD34^−^) are shown. (B) The percentage of peripheral CD34^+^ monocytes (CD3^−^CD14^+^CD16^−^CD34^+^) in total CD14^+^CD16^−^ classical monocytes determined by flow cytometry is shown. The results for cART-untreated HIV-1-infected individuals (“HIV-1+”, n=5) and HIV-1-uninfected donors (“HIV-1−”, n=5) are summarized. \**p* < 0.05. (C) Genomic DNA was isolated from peripheral CD34^+^ monocytes (CD3^−^CD14^+^CD16^−^CD34^+^) and CD34^−^ monocytes (CD3^+^CD14^+^CD16^−^CD34^−^) and subjected to proviral PCR (100 cells/reaction in the first PCR) to amplify *gag* or *pol* region with eight different primer pairs (see Supplemental Table 1). The positive PCR reactions are indicated by orange (for CD34^+^) or cyan (for CD34^−^) in eight squares. The results for cART-untreated HIV-1-infected individuals (“ART−”, n=6) are summarized, which include the results of PBMCs collected from the same individual but different visits (KI110-1 and KI110-2). Plasma viral load (PVL; HIV-1 RNA copies/ml) are also shown. (D) HIV-1 proviral PCR was also performed using PBMCs of individuals treated with cART for more than 3 years (3 – 19 years, median=12.7 years). In the analyses, resting CD4^+^ T cells (CD4^+^CD25^−^CD69^−^HLA-DR^−^), but not CD34^−^ monocytes, were used as a reference. The conditions of PCR were as in (**C**), and the positive PCR reactions are indicated by orange (for CD34^+^ monocytes) or green (for resting CD4^+^ T cells) in eight squares. Individuals (n=16) are grouped (i, ii, iii, and iv) according to the results of positive PCR.

### CD34^+^ monocytes express HIV-1 receptors at higher levels and HIV-1 restriction factors at lower levels than CD34-negative monocytes

To clarify why CD34^+^ monocytes are susceptible to HIV-1, we next analyzed their phenotypes. When cultured with M-CSF, the key cytokine for monocyte-to-macrophage differentiation (19), macrophages derived from CD34^+^ monocytes and CD34-negative monocytes were similar in the expression level of myeloid markers, such as CD14 and CD163, but different in the morphology as CD34^+^ monocyte-derived macrophages were more spindle (Fig. 2A). This result was consistent with the finding that the CD34^+^ fraction contained fibroblastic leukocytes called fibrocytes (12, 17). Indeed, CD34^+^ monocytes and CD34-negative monocytes were overlapping but not necessarily identical in their gene expression as chemokines/chemokine receptors and genes involved in viral replication/viral life cycle were enriched in CD34^+^ monocytes (Fig. 2B, Supplemental Table 2).

**Figure 2.**
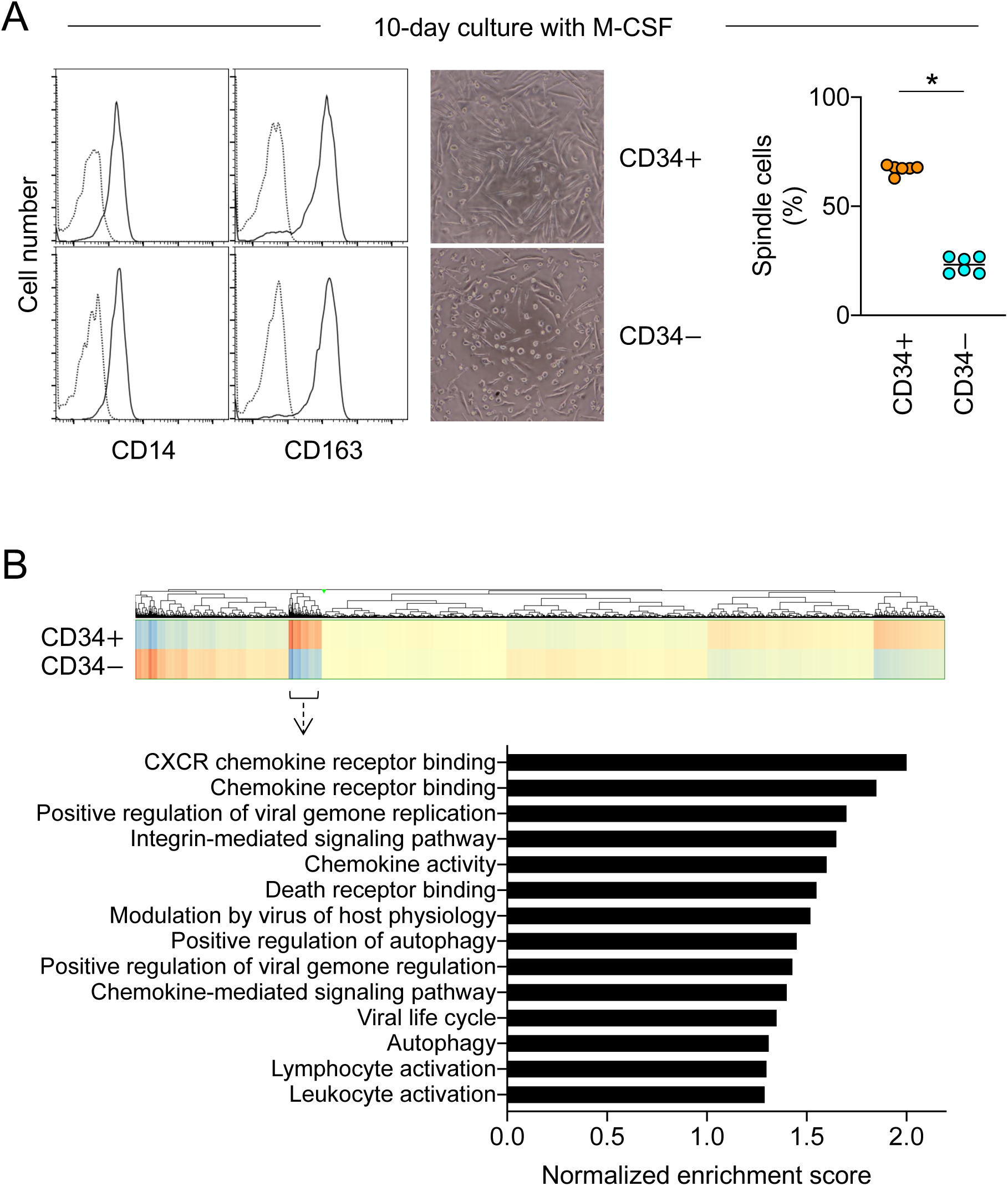
Characteristics of peripheral CD34^+^ monocytes. (A) Peripheral CD34^+^ monocytes (“CD34+”, CD3^−^CD14^+^CD16^−^CD34^+^) and CD34^−^ monocytes (“CD34−”, CD3^+^CD14^+^CD16^−^CD34^−^) of a healthy donor were sorted and cultured in the presence of M-CSF for 10 days. The differentiated macrophages were analyzed for cell surface expression of CD14 and CD163 by flow cytometry (left panels), and morphologies (middle panels). The percentage of spindle cells in those cultures is also summarized (right graph). \**p* < 0.05. (B) Peripheral CD34^+^ monocytes and CD34^−^ monocytes of a healthy donor were sorted and directly subjected to microarray analysis. The pathway enrichment using GSEA method is shown.

In this study, we found that CD34^+^ monocytes expressed HIV-1 receptor CD4 and co-receptor CCR5 at higher levels than CD34-negative monocytes (Fig. 3A, upper panels), which was not observed for other cell surface proteins tested, including CCR2 and MHC-Ⅰ (Fig. 3A, lower panels). Moreover, we found that CD34^+^ monocytes expressed several HIV-1 restriction factors, such as MX2, SAMHD1, and BST2, at lower levels than CD34-negative monocytes (Fig. 3B). Among them, SAMHD1 and MX2 inhibit HIV-1 reverse transcription and HIV-1 nuclear import (23), respectively, both of which are the post-entry early steps in the viral life cycle. Thus, these results suggest that CD34^+^ monocytes readily allow for the entry and reverse transcription of HIV-1, and subsequent viral import into cellular nuclei.

**Figure 3.**
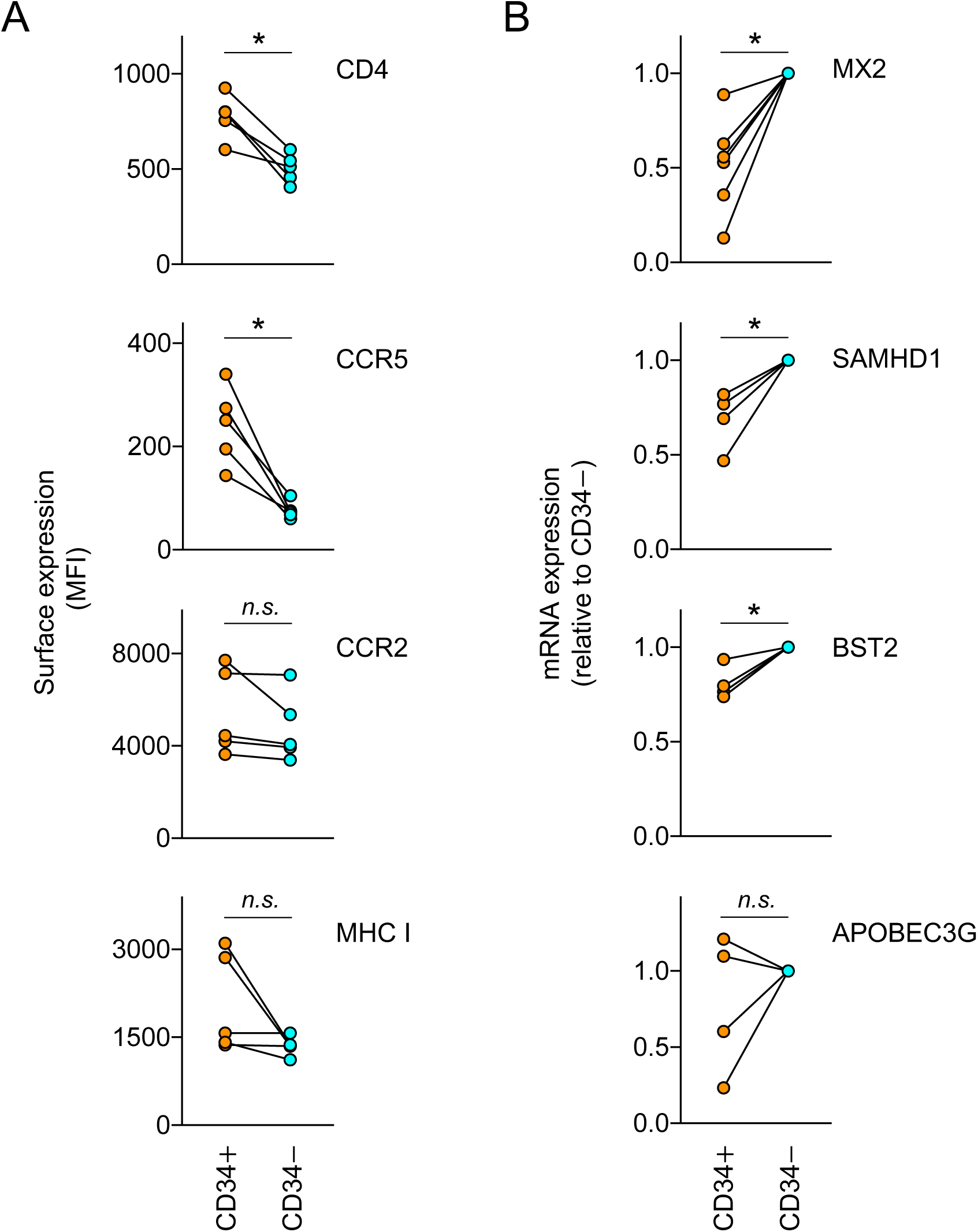
HIV-1-related characteristics of peripheral CD34^+^ monocytes. (A) Peripheral CD34^+^ monocytes (“CD34+”, CD3^−^CD14^+^CD16^−^CD34^+^) and CD34^−^ monocytes (“CD34−”, CD3^+^CD14^+^CD16^−^CD34^−^) of healthy donors (n=5) were analyzed for cell surface expression of CD4, CCR5, CCR2, and HLA-A/B/C (MHC-I) by flow cytometry. The mean fluorescence intensity (MFI) is shown. (B) Peripheral CD34^+^ monocytes and CD34^−^ monocytes of healthy donors (n=4 or 6) were sorted and analyzed for mRNA expression of MX2, SAMHD1, BST2, or APOBEC3G by qRT-PCR. The expression level shown is relative to that of CD34^−^ monocytes. *n.s.*, not significant. \**p* < 0.05.

### CD34^+^ monocytes are present not only in peripheral blood but also lymph nodes, and they share phenotypes including the expression of HIV-1 receptors and regulators of in vivo cellular trafficking

We next attempted to get a clue about how peripheral CD34^+^ monocytes could become infected in individuals whose plasma viral load was below the detection limit by the long-term cART. To this end, we performed experiments using lymph nodes since most HIV-1 reservoirs are located in the tissues (24). Due to limitations of use of tissues of HIV-1-infecetd individuals, we initially analyzed lymph nodes of rhesus macaques infected with the HIV-1-related monkey-tropic virus SIV (19, 20). A substantial part of the CD14^+^ cells was positive for CD34 (Fig. 4A, dot plots). Although its detection frequency on a per cell basis was lower than that of CD4^+^ T cells (Fig. 4A, right panel, green bars), the proviral DNA was detected in the CD34^+^ monocytes-like CD45^+^CD3^−^CD14^+^CD34^+^ cells (Fig. 4A, right panel, orange bar). Consistent with this, the population expressing SIV p27 Gag proteins was detected in the CD34^+^ monocytes-like cells (Fig. 4B). Those infected CD34^+^ monocytes-like cells might escape from the recognition by CD8^+^ cytotoxic T cells since CD14^+^ cells (Fig. 4C, brown) and CD8^+^ T cells (Fig. 4C, green) were present in different regions within lymph nodes.

**Figure 4.**
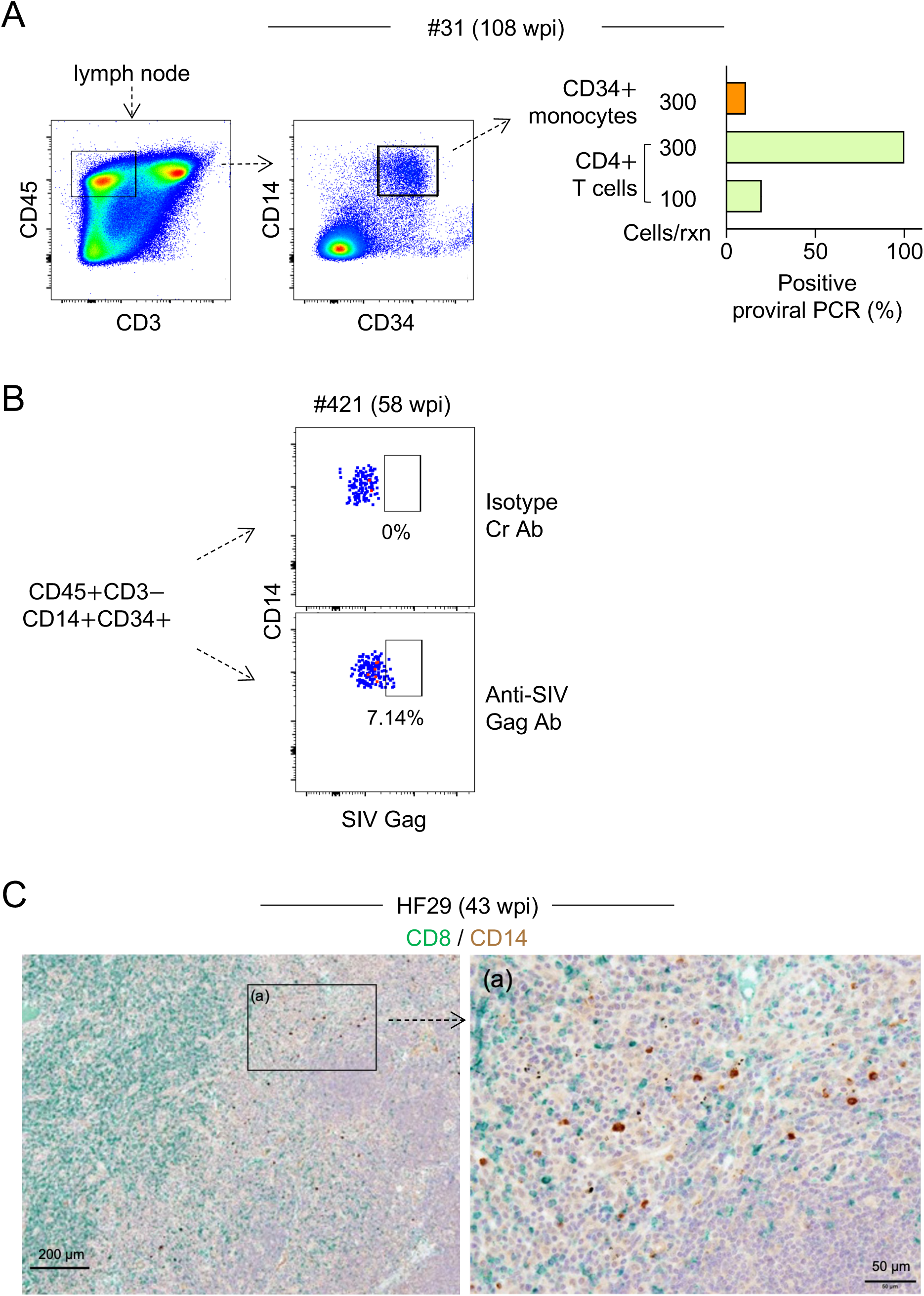
SIV proviral DNA and Gag proteins in lymph node CD34^+^ monocytes of rhesus macaques. (A) In left panels, sorting strategy for CD34^+^ monocytes-like CD45^+^CD3^−^CD14^+^CD34^+^ cells in lymph node of an SIV-infected macaque (#31) (20) is shown. In right graph, the CD34^+^ monocytes and control CD4^+^ T cells (CD45^+^CD3^+^CD8^−^CD19^−^) were subjected to proviral PCR (300 or 100 cells/reaction in the first PCR as indicated). The percentage of the positive PCR reactions are summarized (orange for CD34^+^ monocytes or greenish yellow for CD4^+^ T cells). (B) The CD34^+^ monocytes-like CD45^+^CD3^−^CD14^+^CD34^+^ cells in lymph node of an SIV-infected macaque (#421) (20) were incubated with anti-SIV Gag antibody (lower panel) or isotype control antibody (upper panel), and analyzed for their expression of p27 Gag by flow cytometry. wpi, weeks post-infection. (C) Lymph node of an SIV-infected rhesus macaque (HF29) (19) was analyzed for CD14^+^ cells (brown) or CD8^+^ cells (green) by immunohistochemistry.

To clarify how CD34^+^ monocytes-like cells in lymph nodes relate to those in peripheral blood, we next analyzed lymph nodes resected from five patients with oral squamous cell carcinoma and detected the CD34^+^ monocytes-like CD45^+^CD14^+^CD3^−^CD19^−^CD34^+^ cells (Fig. 5A). In lymph nodes, most CD45^+^CD14^+^CD3^−^CD19^−^ cells expressed the myeloid markers, such as CD163 and CD64 (Fig. 5A, lower right panels), and contained the CD34^+^ fraction (Fig. 4A, upper panel). Thus, we next analyzed whether the CD34+monocytes-like cells in lymph nodes were identical to those in peripheral blood. As previously reported (17), we confirmed that CD34^+^ monocytes in peripheral blood expressed a cell surface molecule signaling lymphocytic activation molecule family member 7 (SLAMF7) at higher levels than CD34-negative monocytes (Fig. 5B, right panel). The CD34^+^ monocytes-like cells in lymph nodes also expressed SLAMF7 at higher levels than the CD34-negative fraction (Fig. 5B, left panel). Importantly, the CD34^+^ monocytes-like cells in lymph nodes also expressed CD4 and CCR5, but not CCR2, at higher levels than the CD34-negative fraction (Fig. 5C), as observed for circulating CD34^+^ monocytes (see Fig. 3A). Finally, we found that both lymph node CD34^+^ monocytes-like cells (Fig. 5D, left panels) and circulating CD34^+^ monocytes (Fig. 5D, right panels) highly expressed CCR7 and sphingosine-1-phosphate receptor 1 (S1PR1), both of which are critical regulators of in vivo cellular trafficking (25–33). These results suggest that CD34^+^ monocytes have a high potential of in vivo trafficking and are therefore detectable in both peripheral blood and lymph nodes.

**Figure 5.**
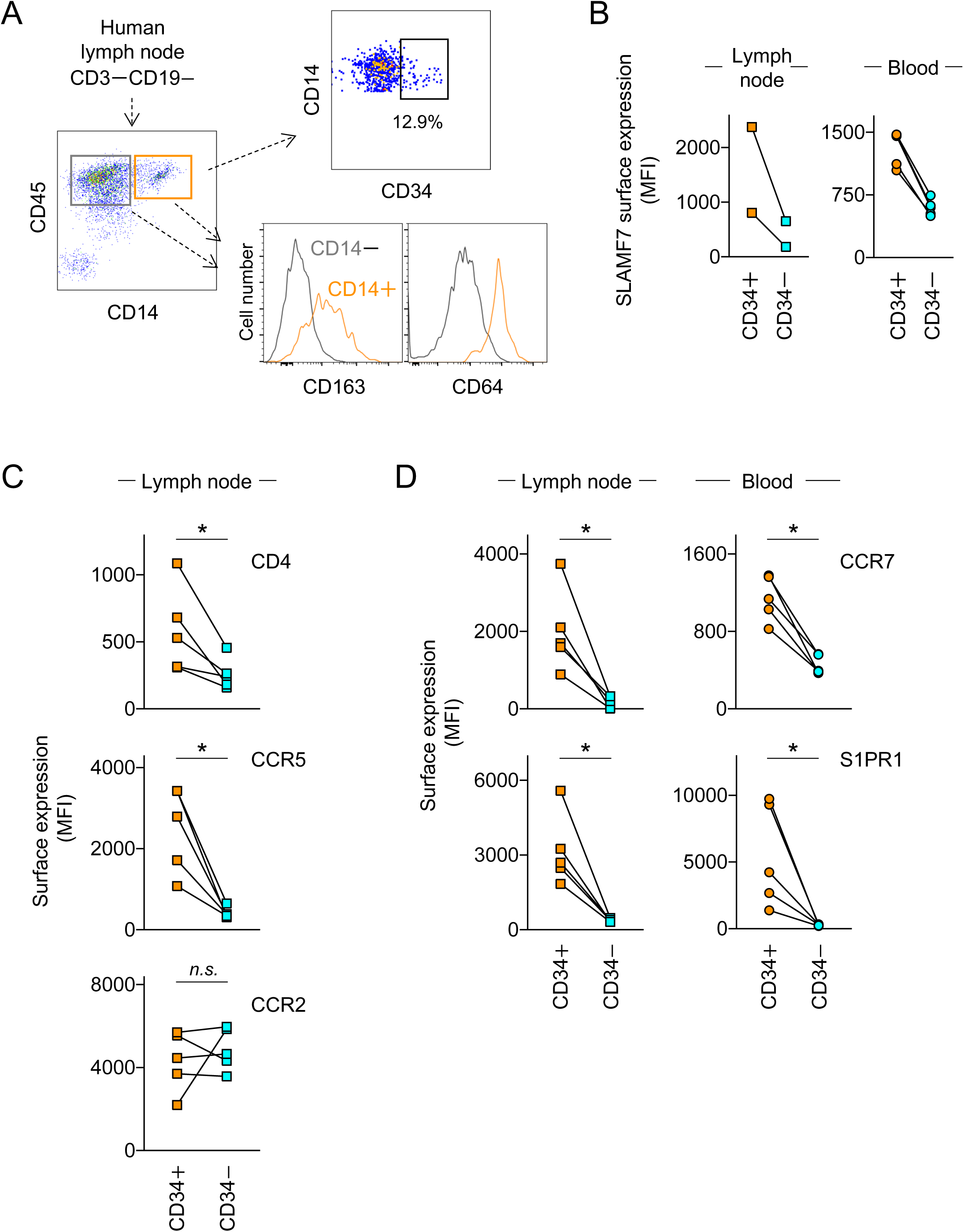
Characteristics of lymph node CD34^+^ monocytes. (A) Typical example of flow cytometric analysis of lymph nodes obtained from patients with oral squamous cell carcinoma is shown. The CD3^−^CD19^−^ fraction was analyzed for cell surface expression of CD45 and CD14 (left panel). The two fractions (CD45^+^CD14^−^ and CD45^+^CD14^+^) were further analyzed for cell surface expression of CD163 and CD64 (lower right panels). The CD45^+^CD14^+^ fraction was also analyzed for cell surface expression of CD34 (upper right panel). (B) The CD34^+^ monocyte fraction and CD34^−^ monocyte fraction in lymph nodes obtained from patients with oral squamous cell carcinoma (see the upper right panel of panel **A**) were further analyzed for cell surface expression of SLAMF7 (left panel, n=2). Peripheral CD34^+^ monocytes and CD34^−^ monocytes obtained from healthy volunteers were also analyzed as a reference (right panel, n=5). The mean fluorescence intensity (MFI) is shown. (C) The CD34^+^ monocyte fraction and CD34^−^ monocyte fraction in lymph nodes obtained from patients with oral squamous cell carcinoma were analyzed for cell surface expression of CD4, CCR5, and CCR2 (n=5). MFI is shown. (D) The CD34^+^ monocyte fraction and CD34^−^ monocyte fraction in lymph nodes obtained from patients with oral squamous cell carcinoma were analyzed for cell surface expression of CCR7 and S1PR1 (left panels, n=5). Peripheral CD34^+^ monocytes and CD34^−^ monocytes obtained from healthy volunteers were also analyzed as a reference (right panels, n=5). MFI is shown. *n.s.*, not significant. \**p* < 0.05.

## Discussion

We previously reported that HIV-1 proviral DNA was enriched in the CD34^+^ fraction within classical monocytes of cART-untreated individuals (17). Although the underlying mechanisms have been unclear, the high susceptibility of CD34^+^ monocytes to HIV-1 can be explained by their high expression of HIV-1 receptor/co-receptor, and low expression of host factors that restrict the post-entry early steps of HIV-1 life cycle, such as MX2 and SAMHD1, as shown in this study (Fig. 3). A noteworthy finding of this study was the proviral persistence in CD34^+^ monocytes of individuals on cART (Fig. 1D). Thus, this study identified CD34^+^ monocytes as a novel candidate of long-lived latently or persistently infected monocytes.

We revealed that CD34^+^ monocytes are present in both peripheral blood and lymph nodes (Figs. 4, 5), and highly express CCR7 and S1PR1 (Fig. 5). CCR7 and S1PR1 are critical regulators of in vivo cellular trafficking (25–33). For instance, CCR7^-/-^ mice had a decreased number of monocytes in lymph nodes during *Toxoplasma* infection (27), and CCR7-deficient LPS-primed monocytes failed to migrate from footpads to lymph nodes (28). Consistent with this, forced CCR7 expression guided injected monocytes to lymph nodes (33). The role of S1PR1 in guiding cells from tissues to circulation is also well known (32). For instance, S1PR1-acting FTY720 (also known as Fingolimod) which blocks cellular exit from lymph nodes (34), reduced circulating monocytes (29) and the lymph node-to-lymph node trafficking of *yersinia pestis*-infected monocytes (31). Thus, it is likely that CD34^+^ monocytes circulate between peripheral blood and tissues including lymph nodes more potently than classical monocytes. The potent proliferative capacity (13, 15) and trans-endothelial migration/reverse trans-endothelial migration capacity (35, 36) of CD34^+^ monocytes may further support their circulation between peripheral blood and tissues. Taken together, the present study raises an unreported possibility that CD34^+^ monocytes are infected with residual HIV-1 in lymph nodes of individuals on cART owing to their high susceptibility to HIV-1, and return to circulation owing to their potent trafficking characteristics. The idea can explain the presence of proviral DNA in peripheral CD34^+^ monocytes even after long-term cART.

It has been shown that intermediate CD14^+^CD16^+^ monocytes, the monocyte subset distinct from CD34^+^ monocytes in classical monocytes, are also susceptible to HIV-1 (37, 38). CD14^+^CD16^+^ monocytes expressed HIV-1 co-receptor CCR5 more highly than total classical monocytes, and harbored HIV-1 proviral DNA more frequently than total classical monocytes in individuals on cART (37). These features may resemble CD34^+^ monocytes, but it is unclear how CD14^+^CD16^+^ monocytes express CCR7 and S1PR1 and whether they are present in lymph nodes. Interestingly, an in vivo 6,6-^2^H_2_-glucose labeling experiment using healthy volunteers and eosinophilic asthma patients proposed the model that CD14^+^CD16^+^ monocytes stay outside for 1.6 days and return to the circulation (39), although it is unclear whether CD14^+^CD16^+^ monocytes localize to lymph nodes before recirculation. Thus, it is possible that the monocyte population harboring Gag DNA (135.7 copies/million cells) that Veenhuis *et al.* recently identified in individuals on cART (7) is consisted of CD34^+^ monocytes and CD14^+^CD16^+^ monocytes. To test this hypothesis, the virological and phenotypical features of these two monocyte populations should be compared in detail under the same setting.

Due to the limitation of volumes of peripheral blood collected and relative low number of monocytes in PBMCs, it remains unexplored to what extent CD34^+^ monocytes and CD14^+^CD16^+^ monocytes serve as HIV-1 reservoir that produces infections virions. Meanwhile, cells harboring defective proviruses are another important target towards HIV-1 cure since defective or deleted proviruses are not necessarily transcriptionally silent but can be produce viral proteins such as Nef and Gag (40–42), which could alter anti-viral T cell response and myeloid cell function that contribute to HIV-1-associated chronic comorbidities (8). Thus, it would be intriguing to test how CD34^+^ monocytes and CD14^+^CD16^+^ monocytes express viral mRNAs and proteins.

CD4^+^ T cells are the major HIV-1 reservoir, but recent studies suggest that myeloid cells also meet the definition of a clinically relevant reservoir (4–8). The present study, in which we identified CD34^+^ monocytes as the second candidate of proviral-persistent monocyte subset, will be helpful for the elimination of myeloid reservoir.

## Acknowledgments

We thank K. Tomoda, K. Nasu, and I. Suzu for their secretarial and technical assistance. We also thank Editage for the English language editing and review.

