## Supplemental Tables for "CD34-positive monocytes are highly susceptible to HIV-1"

Supplemental Table 1. PCR primers

| Region | PCR | Sequence |
|---|---|---|
| 1 | 1st | 5'-CAGACCCTTTTAGTCAGTGTGGAAAAT[illegible] |
| | | 5'-ACAATCTTTCATTTGGTGTCCTTC-3' |
| | 2nd | 5'-CTAGCAGTGGCGCCCGAACA-3' |
| | | 5'-TTTCCACATTTCCAACAGCC-3' |
| 2 | 1st | 5'-GAGCCACCCCACAAGATTTA-3' |
| | | 5'-ACAATCTTTCATTTGGTGTCCTTC-3' |
| | 2nd | 5'-GACAAGGACCAAAGGAACCC-3' |
| | | 5'-TTTCCACATTTCCAACAGCC-3' |
| 3 | 1st | 5'-GACAAGGACCAAAGGAACCC-3' |
| | | 5'-ACAATCTTTCATTTGGTGTCCTTC-3' |
| | 2nd | 5'-TAGAAGAAATGATGACAGCATG-3' |
| | | 5'-TTTCCACATTTCCAACAGCC-3' |
| 4 | 1st | 5'-TTTTTGACTAGCGGAGGCTAGAA-3' |
| | | 5'-ACAATCTTTCATTTGGTGTCCTTC-3' |
| | 2nd | 5'-TTTTTGACTAGCGGAGGCTAGAA-3' |
| | | 5'-TTTCCACATTTCCAACAGCC-3' |
| 5 | 1st | 5'-CAGGAATTTGGCATTCCCTA-3' |
| | | 5'-TGGAGACTCCCTGACCCAAATG-3' |
| | 2nd | 5'-ATTGGGGGGTACAGTGCAG-3' |
| | | 5'-TGGAGACTCCCTGACCCAAATG-3' |
| 6 | 1st | 5'-AAAAGGGCTGTTGGAAATGT-3' |
| | | 5'-GTATTGACAAACTCCCACTCAG-3' |
| | 2nd | 5'-ACAGGCTAATTTTTTAGGGAA-3' |
| | | 5'-TCATAATACACTCCATGTAC-3' |
| 7 | 1st | 5'-CAGGAATTTGGCATTCCCTA-3' |
| | | 5'-CTAATCCTCATCCTGTCTAC-3' |
| | 2nd | 5'-TCCACTTTGGAAAGGACCAG-3' |
| | | 5'-CTAATCCTCATCCTGTCTAC-3' |
| 8 | 1st | 5'-ACACCTGTCAACATAATTGG-3' |
| | | 5'-TGTATGTCATTGACAGTCCA-3' |
| | 2nd | 5'-GGGCCTGAAAATCCATACAA-3' |
| | | 5'-GGTGATCCTTTCCATCCCTG-3' |

Supplemental Table 2. Genes enriched in $CD34^{+}$ monocytes (>two-fold)

| Chemokine/chemokine receptor-related | Viral replication/viral life cycle-related |
|---|---|
| CCL5 | ITGB3 |
| CXCL10 | ATG7 |
| CXCL11 | TFRC |
| CXCL9 | RANBP2 |
| CCR2 | NFIA |
| CXCL1 | ZNF639 |
| CCL2 | ITGB5 |
| CX3CR1 | SLC10A1 |
| CXCL2 | TBC1D20 |
| CCR5 | NUP88 |
| CXCL5 | RPS24 |
| CXCR5 | WWP2 |
| CXCR4 | KPNA1 |
| CCL18 | ATP6V0E2 |
| CXCL12 | CR1 |
| CCL16 | VTA1 |
| CCL28 | SEH1L |
| CCL3 | ATG5 |
| CX3CL1 | NEDD4L |
| CXCR3 | IDE |
| CCL23 | CD209 |
| CXCL16 | |
| CXCL14 | |
| CCL24 | |
| CXCR2 | |
| CCL20 | |
| CCR6 | |
| CCL11 | |
| CXCL8 | |
| CCL1 | |
| CXCL3 | |
